# Professional medical writing support and the quality, ethics and timeliness of clinical trial reporting: a systematic review

**DOI:** 10.1101/501403

**Authors:** Obaro Evuarherhe, William Gattrell, Richard White, Christopher C Winchester

## Abstract

**Background:** Many authors choose to work with professional medical writers when reporting the results of clinical trials. We conducted a systematic review to examine the relationship between professional medical writing support (PMWS) and the quality, ethics and timeliness of publications reporting clinical trials.

**Methods:** Using terms related to ‘medical writer’ and ‘observational study’, we searched MEDLINE and Embase (no date limits), as well as abstracts and posters from meetings of the International Society for Medical Publication Professionals (ISMPP; 2014–2017). We also hand-searched the journals *Medical Writing* and *The Write Stuff* (2014–2017), and the bibliographies of studies identified in the electronic searches. We screened the results to identify studies that compared the quality, ethics and timeliness of clinical trial publications written with and without declared PMWS.

**Results:** Our searches identified 97 potentially relevant studies, of which 89 were excluded during screening and full paper review. The remaining eight studies compared 849 publications with PMWS with 2073 articles developed without such support. In these eight studies, PMWS was shown to be associated with: increased adherence to Consolidated Standards of Reporting Trials (CONSORT) guidelines (in 3/3 studies in which this was assessed); publication in journals with an impact factor (one study); a higher quality of written English (one study); and a lower likelihood of reporting non-pre-specified outcomes (one study). PMWS was not associated with increased adherence to CONSORT for Abstracts guidelines (one study) or with the impact of published articles (mean number of citations per year, mean number of article views per year and Altmetric score; one study). In studies that assessed timeliness of publication, PMWS was associated with a reduced time from last patient visit in clinical trials to primary publication (one study), whereas time from submission to acceptance showed inconsistent results (two studies).

**Conclusions:** This systematic review of eight observational studies suggests that PMWS increases the overall quality of reporting of clinical trials and may improve the timeliness of publication.

## Background

Timely and complete reporting of the results of clinical trials is an ethical imperative [1]; it helps to eliminate duplicative effort, enables researchers to develop more up-to-date study hypotheses and allows clinicians and patients to judge the benefits or risks of different therapies. Although the pharmaceutical industry has made great strides to address criticism for a perceived lack of transparency in the disclosure of clinical trial results, the quality, ethics and timeliness of clinical trial reporting remain closely scrutinized for both industry-funded and academically funded trials [2–6].

Pharmaceutical companies often offer authors professional medical writing support (PMWS) to assist in the reporting of clinical trial results [7]. International guidelines endorse the acknowledgement of PMWS [8,9], and the proportion of articles in the medical literature with such an acknowledgement is 6–19% [7,10,11]. We conducted a systematic review to identify and analyse published studies that investigated the association between PMWS and the quality, ethics and timeliness of clinical trial reporting.

## Methods

### Systematic literature search

Published studies relating to medical writing were identified through a systematic literature review. Cochrane, Embase, MEDLINE In-Process & Other Non-Indexed Citations, and MEDLINE 1946–contacted the corresponding present were searched on 8 March 2018 via the Ovid platform.

The search strategy comprised terms relating to medical writing, medical publication professional and medical communication, and was combined with terms for observational, cross-sectional or epidemiological studies, with no limits on date, language or country in which the research was conducted (Figure 1).

**Figure 1.**
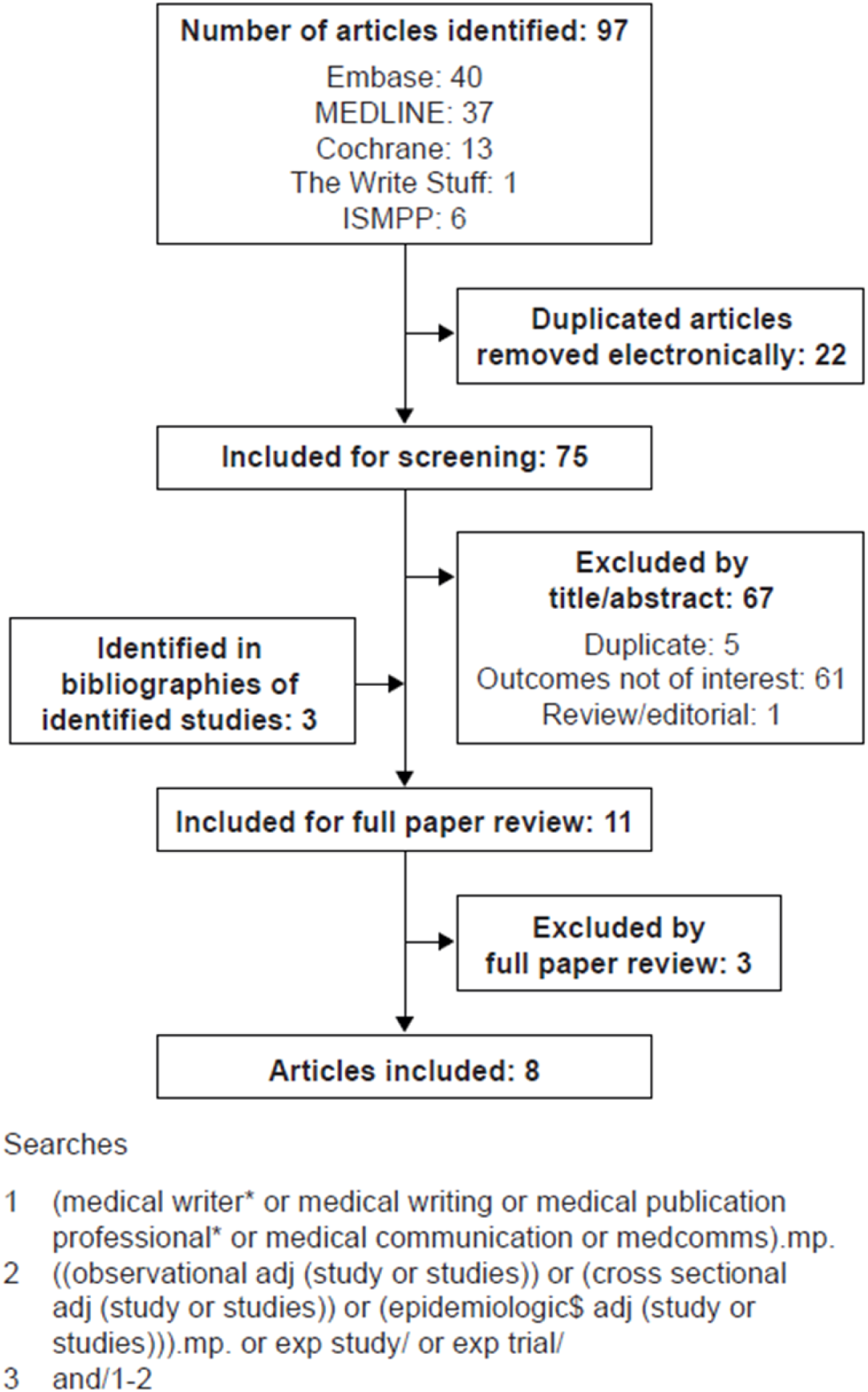
PRISMA diagram of included and excluded studies. ISMPP, International Society for Medical Publication Professionals; PRISMA, Preferred Reporting Items for Systematic Reviews and Meta-Analyses.

### Supplementary searches

Supplementary searches were conducted of the International Society for Medical Publication Professionals (ISMPP) congress proceedings (which are published as supplementary articles in *Current Medical Research Opinion*), and the journals *Medical Writing* and *The Write Stuff* (which are available via the European Medical Writers Association [EMWA] website) using the terms ‘medical writ*’ and ‘medical publication professional’. Supplementary searches were limited to 2014–2017. We contacted the corresponding authors of congress abstracts identified in the supplementary searches to request access to full posters/presentations. The bibliographies of studies identified in the electronic searches were also reviewed to identify additional relevant references.

### Study selection and data collection

All identified studies were screened against inclusion and exclusion criteria in accordance with the 2009 Preferred Reporting Items for Systematic Reviews and Meta-Analyses (PRISMA) guidelines [12]. For congress abstracts identified in the supplementary searches, full posters were obtained from the ISMPP website or from the authors. Identified congress abstracts were excluded as ‘duplicates’ if a full version of the study had been published. Studies eligible for inclusion were in English, and compared the quality, ethics or timeliness of articles reporting clinical trials that had been developed with and without acknowledged PMWS. Studies that did not directly compare clinical trial publications that had been developed with and without PMWS were excluded, as were those that reported outcomes that were unrelated to quality, ethics or timeliness, and those that assessed study types other than clinical trials.

Details of study methodology, study size, main outcome measures, quality-related outcomes (e.g. adherence to Consolidated Standards of Reporting Trials [CONSORT] or CONSORT for Abstracts [CONSORT-A]), ethics-related outcomes (e.g. reporting of non-pre-specified outcomes) and timeliness-related outcomes were extracted from each eligible study. The influence of PMWS was classified as positive, non-significant or negative for each study, based on the results and statistical analyses reported in each publication.

## Results

### Search results

Our searches identified 75 potentially relevant publications after exclusion of 22 duplicate publications; 70 were excluded during screening and full paper review, and three were identified in bibliographies of identified studies (Figure 1). Of the eight included studies, three were full publications (two in peer-reviewed journals [13,14], one in a non-peer reviewed journal [15]), and five were congress abstracts (four poster presentations [16–19], one oral presentation [20]). Although no date limit was included in the search strategy, only two of the identified studies were published before 2015: one in 2006 [7] and the other in 2010 [15] (Table 1). The eight included studies analysed 849 articles that had been developed with PMWS and 2073 articles developed without.

**Table 1.** Overview of included studies

| First author, year | Number of included studies |  | Publication type | Description of analysed articles |
| --- | --- | --- | --- | --- |
|  | With PMWS | Without PMWS |  |  |
| Gattrell, 2016 [14] | 110 | 123 | Peer-reviewed publication | Articles reporting RCT results published in BioMed Central journals<br>Biomed Central journals have been used in previous studies of adherence to CONSORT guidelines [37] |
| Gattrell, 2016 [17] | 110 | 123 | Poster presentation | Articles reporting RCT results published in BioMed Central journals (same cohort of articles analysed in Gattrell *et al.* [14]) |
| Gattrell, 2017 [16] | 17 | 49 | Poster presentation | Sub-analysis of outcomes reported in the top five medical journals comparing each article with its corresponding study protocol or clinical trial registry entry using publicly available COMPare data<br>The COMPare project is evaluating outcome reporting in clinical trials by comparing publications with the respective registry entries [22] |
| Jacobs, 2010 [15] | 152 | 69 | Non-peer-reviewed publication | RCTs published between October 2004 and August 2009 in the journal *Current Medical Research and Opinion* |
|  |  |  |  | • *Current Medical Research and Opinion* almost exclusively publishes industry-funded studies |
| Mills, 2017 [13] | 66 | 397 | Peer-reviewed publication | • RCTs published between 2011 and 2014 in five high-impact medical journals: *The New England Journal of Medicine*, *Annals of Internal Medicine*, *The Lancet*, *The BMJ* and *JAMA*<br>• All included articles had been analysed in a cross-sectional study examining reporting quality of RCTs [38] |
| Shah, 2015 [19] | 40 | 103 | Poster presentation | • Neuroscience and cardiology RCTs published between 2009 and 2014 in different journals from the Asia-Pacific region |
| Shah, 2016 [18] | 404 | 392 | Poster presentation | • RCTs conducted to gain US FDA approval in 2014<br>• Innovative and novel drugs and new molecules approved by the FDA in 2014, identified in FDA reports |
| Woolley, 2006 [7] | 60 | 940 | Congress abstract | • Original research articles published up to January 2005 from each of 10 high-impact factor, international, peer-reviewed medical journals from a range of therapeutic areas |

COMPare, Centre for Evidence-Based Medicine Outcome Monitoring Project; CONSORT, Consolidated Standards of Reporting Trials; FDA, Food and Drug
Administration; PMWS, professional medical writing support; RCT, randomized controlled trial.

### Quality of reporting

Of the identified studies comparing articles developed with and without PMWS, three assessed adherence to CONSORT guidelines [14,15,19]. Each of these studies, using a different statistical approach to assess adherence, showed that PMWS was associated with increased adherence to CONSORT guidelines (Table 2). Articles developed with PMWS were significantly more likely to report completely at least 50% of the assessed CONSORT items (*p* < 0.05) [14,21] and to comply with more CONSORT items than articles without PMWS (*p* < 0.05) [15]. Similarly, articles with 80–100% compliance with CONSORT items were significantly more likely to have been developed with PMWS than those with less than 80% compliance (*p* < 0.0001) [19]. Looking at individual CONSORT items, one identified study showed that articles with PMWS were significantly more likely to report all important adverse events or side effects than those without PMWS [15], and another showed that PMWS increased adherence to six of 12 CONSORT items assessed: specification of primary outcome; sample size calculation; type of randomization; publication of a participant flow diagram; provision of dates defining recruitment and follow-up; and details of trial registration [14]. Additionally, in this study, another CONSORT item (who generated the allocation sequence) was only reported in 5/110 articles developed with PMWS and none of the 123 articles without PMWS; thus, a relative risk could not be calculated [14]. One additional study assessed adherence to CONSORT-A and showed that PMWS was not associated with an overall increase in adherence [13]; PMWS was associated with lower levels of adherence with respect to reporting of study setting and higher levels of adherence in relation to disclosure of harms/side effects and funding source in the abstract [13].

**Table 2.** Summary of results

| First author, year | Outcome measured | Effect of PMWS |  |  |
| --- | --- | --- | --- | --- |
|  |  | Positive | Non-significant | Negative |
| Gattrell, 2016 [14] | Adherence to CONSORT guidelines | The proportion of articles that completely reported at least 50% of the assessed CONSORT items<br>• With PMWS: 43/110 articles (39.1%; 95% CI: 29.9–48.9)<br>• Without PMWS: 26/123 articles (21.1%; 95% CI: 14.3–29.4) |  |  |
| Jacobs, 2010 [15] |  | Logistic regression analysis showed that CONSORT items were significantly more likely to be completed in papers with a clear acknowledgement of PMWS than in those without<br>(OR 1.44; 95% CI: 1.04–2.00; $p = 0.03$) |  |  |
| Shah, 2015 [19] |  | 23/97 articles with PMWS (24%) had |  |  |
| | | 80–100% CONSORT adherence, whereas 5/105 articles developed without PMWS (5%) had 80–100% CONSORT adherence ($p < 0.0001$) | | |
| Mills, 2017 [13] | Adherence to CONSORT-A guidelines |  | The mean proportion of CONSORT-A items reported was similar with and without PMWS (64.3% vs 66.5%, respectively; $p = 0.30$); PMWS was associated with a lower level of compliance with reporting of study setting (RR 0.40; 95% CI: 0.23–0.70) and a higher level of adherence to disclosure of harms/side effects (RR 2.04; 95% CI: 1.37–3.03) and funding source (RR 1.75; 95% CI: 1.18–2.60) |  |
| Gattrell, 2016 [14] | Quality of written English | Proportion of articles rated by all reviewers during peer review as having an acceptable standard of written English<br>• With PMWS: 81.1% (43/53 articles; 95% CI: 67.6–90.1)<br>• Without PMWS: 47.9% (23/48 articles; 95% CI: 33.5–62.7) |  |  |
| Gattrell, 2016 [17] | Publication in journal with an impact factor | Likelihood of publication in journal with an impact factor was significantly improved with PMWS ($p = 0.001$) | | |
| | Mean impact factor of publication | Mean impact factor of publication was significantly improved with PMWS ($p < 0.001$) | | |
| Gattrell, 2017 [16] | Reporting of non-pre-specified outcomes | Articles developed with PMWS reported fewer non-pre-specified outcomes than both industry-funded ($p = 0.028$) and non-industry-funded articles ($p < 0.01$) developed without PMWS | | |
| Gattrell, 2016 [17] | Mean number of citations per year | | Mean number of citations per year was not significantly improved with PMWS ($p = 0.11$) | |
| | Mean number of article views per year | | Mean number of article views per year was not significantly improved with PMWS ($p = 0.84$) | |
| | Altmetric score | | Altmetric score was not significantly improved with PMWS ($p = 0.55$) | |
| Gattrell, 2016<br>[14,21] | Manuscript acceptance<br>time | | | Time from manuscript submission to acceptance was increased with PMWS (167 days [IQR 114.5–231 days] vs 136 days [IQR 77–193 days], $p < 0.01$); mean number of versions submitted was unchanged |
| Shah, 2016 [18] | Time to publication | Time to publication from last patient visit in clinical trials was reduced with PMWS (18.6 [SD 13.2] months vs 30.8 [SD 11.7] months) |  |  |
| Woolley, 2006 [7] | Manuscript acceptance<br>time | | Time from manuscript submission to acceptance was reduced with PMWS (83.6 days vs 132.2 days), although this difference was not statistically significant ($p = 0.053$) | |

CI, confidence interval; CONSORT, Consolidated Standards of Reporting Trials; CONSORT-A, CONSORT for Abstracts; IQR, interquartile range; OR, odds ratio;
PMWS, professional medical writing support; RR, relative risk; SD, standard deviation.

Two studies which represented different analyses of the same group of articles looked at other markers of quality in reporting (Table 2) [14,17]. In these studies, PMWS was positively associated with various measures of reporting quality, including a higher standard of written English (*p* < 0.01) [14,21], higher likelihood of publication in a journal with an impact factor (*p* = 0.001) [17], and higher mean impact factor of the journal accepting the article (*p* < 0.001) [17]. However, there was no association between PMWS and article-level measures of impact, such as mean number of citations per year (*p* = 0.11), mean number of article views per year (*p* = 0.84) and Altmetric score (*p* = 0.55) (Table 2) [17].

### Ethics of publication

Of the identified studies, one examined the relationship between outcome reporting and PMWS using data from the publicly available Centre for Evidence-Based Medicine Outcome Monitoring Project (COMPare) [22]. PMWS was associated with the reporting of fewer non-pre-specified outcomes (*p* = 0.028) [16].

### Timeliness of publication

Three studies looked at the timeliness of clinical trial reporting in articles developed with or without PMWS (Table 2) [14,18,20]. The only study investigating the complete manuscript development time, from last patient visit in clinical trials to article publication, showed that PMWS was associated with reduced time to publication [18]. Two studies investigating the timing of one step in the process, from manuscript submission to acceptance, showed inconsistent results [14,20]. In the first of these studies, PMWS was associated with increased time from manuscript submission to acceptance, although the mean number of versions submitted was unchanged [14]; in the second study, time from manuscript submission to acceptance was reduced, but not significantly [20].

## Conclusions

This systematic review aimed to identify and evaluate studies assessing the effects of PMWS on quality, ethics and timeliness of clinical trial reporting. Overall findings from eight studies assessing 849 articles developed with PMWS and 2073 articles developed without PMWS suggest a positive association between PMWS and improvements in clinical trial reporting. These results were consistent across measures of quality (adherence to CONSORT guidelines and quality of written English), ethics (reporting of non-pre-specified outcomes) and timeliness (time to publication). The improvement in CONSORT adherence associated with PMWS is perhaps unsurprising, given that professional medical writers are routinely trained in Good Publication Practice (GPP3) for the development of peer-reviewed manuscripts [23]; GPP3 guidelines state that authors should follow established reporting standards, including CONSORT [8,9]. Although PMWS was associated with improved adherence to CONSORT, it was not associated with improved adherence to CONSORT-A, suggesting that although professional medical writers improve disclosure overall, they may need to prioritize improving the reporting in the abstract (which is all that is read by many readers).

The improvements in manuscript quality may not be reflected by increased article impact and social media attention. In the one study identified in our systematic review, which examined measures of article impact, there were no significant differences between articles developed with and without PMWS in relation to Altmetric score, number of citations per year and number of article views per year. Medical publications professions have no influence on the subject matter or relevance of the clinical trial and, as such, PMWS may not be expected to affect an article’s post-publication impact.

It is important that authors remain transparent about which specific clinical trial outcomes will be measured and reported. The COMPare project determined the proportion of pre-specified and non-pre-specified outcomes that were reported in clinical studies published in the top five medical journals over a 3-month period [22]. In the present systematic review, one included study conducted a sub-analysis of the publicly available COMPare data and assessed the relationship between PMWS and outcome reporting. The authors found that fewer non-pre-specified outcomes were reported for articles developed with PMWS than for those developed without. This is not the only study to have shown a positive association between PMWS and publication ethics. For instance, a recent study showed that PMWS is associated with increased transparency relating to the source of funding, the author disclosures of financial interest and the acknowledgements of conflicts of interest (or lack thereof) in health economics and outcomes research publications [24]; another study showed that, of 214 publications retracted owing to misconduct between January 1966 and February 2008, only three declared PMWS [25].

One included study looking at the influence of PMWS on timeliness found that PMWS was associated with reduced time from last patient visit to article publication. This period includes processes in which professional medical writers are involved and have a major role, namely manuscript preparation, editing and submission. Two other included studies that examined the influence of PMWS on time from manuscript submission to acceptance revealed mixed results. One of the studies found that time to acceptance was reduced with PMWS, but that the difference was not statistically significant. The other study found that time to acceptance was increased with PMWS; however, it should be noted that the period from submission to article acceptance is not primarily the responsibility of professional medical writers.

Clinicians have reported lack of time as a common reason for non-publication of research findings [26–28]. By specializing in preparation of clinical trial publications, professional medical writers are well placed to aid in the rapid dissemination of trial findings under the direction of the authors, subject to strict publication guidelines [9]. In fact, results from a recent survey showed that authors who use PMWS were more likely to have published as first author at least once in the previous 2 years [29], suggesting that PMWS can also improve overall publication rates.

This systematic review has some limitations, notably that study inclusion was largely based on the assumption that differences in outcomes were attributable to PMWS. It is possible that other factors caused these differences in quality and timeliness. This issue may affect the results of individual studies, but this systematic review combined results from different studies looking at different outcomes of interest, and showed a consistent benefit of PMWS on manuscript quality (including adherence to publication guidelines, quality of written English and publication in high-quality journals), ethics (reporting of pre-specified outcomes) and timeliness (time from completion of trial to publication). Taken together, the findings of this systematic review support the conclusion that PMWS has a positive impact on the high-quality, ethical and timely dissemination of clinical trial data.

The included studies classified articles as having been developed with PMWS only when there was a clear acknowledgement of this support. As such, it is possible that some of the studies classified as having been developed with no PMWS might have had PMWS but had simply failed to acknowledge it. By classifying publications with no clear acknowledgment of PMWS as ‘without PMWS’, the studies identified in this systematic review may have underestimated the effects of PMWS. To minimize the risk of publication bias we employed a broad search strategy with no limits on date, country, language or type of observational study. Most of the identified studies were sourced from conference proceedings (for which the full poster or oral presentation was available in 4/5 cases) and one was published in a non-peer-reviewed journal.

In the identified studies, the outcome measures chosen were widely accepted as measures of quality and completeness. For instance, CONSORT is an independently developed measure of reporting standards recommended by the International Committee of Medical Journal Editors and also medical publications and medical writing societies, including ISMPP, EMWA and the American Medical Writers Association [9]. Other outcomes of interest assessed in this review were assigned independently of the investigators involved in each of the articles analysed in each included study (e.g. standard of written English – assessed during peer review of analysed articles [17]). As such, in this systematic review, we have been successful in analysing a range of outcomes assessed in observational (‘real-world’) studies in a standardized manner that minimizes publication bias.

Further research is needed to elucidate the role of PMWS in clinical trial publication, particularly with regard to productivity and added value [30]. Further research is also required to assess the impact of PMWS in other types of studies published by the pharmaceutical industry, such as observational studies and systematic reviews. As our systematic review identified that most studies of PMWS have only been presented at conferences or published in non-peer-reviewed journals, it is crucial that future studies are published in full in peer-reviewed journals.[31]

Currently, the pharmaceutical industry is more likely than non-industry institutions to disclose clinical trial results properly [32]. This is probably due to a larger investment in internal processes and infrastructure, which includes the use of professional medical writing support. In fact, there have been calls for professional medical writers and publication experts to be employed by academic institutions [33,34]. Additionally, in a survey looking at attitudes to PMWS, academic and clinician respondents to an online survey were generally accepting of PMWS, particularly its influence on editing, journal styling and adherence to reporting guidelines, with 84% of respondents stating that they valued PMWS [35]. In this survey, 82.9% of respondents felt that it was acceptable to receive PMWS; in another survey, PMWS was seen as ‘adding value to publication development’ by almost 90% of participants [35]. Our systematic review appraising current research in this area helps to substantiate the positive attitude to PMWS that is held by clinical and academic professionals seeking to ensure the ethical, accurate and timely publication of clinical trials.

**List of abbreviations**
CI: confidence interval
COMPare: Centre for Evidence-Based Medicine Outcome Monitoring Project
CONSORT: Consolidated Standards of Reporting Trials
CONSORT-A: CONSORT for Abstracts
EMWA: European Medical Writers Association
FDA: Food and Drug Administration
GPP: Good Publication Practice
IQR: interquartile range
ISMPP: International Society for Medical Publication Professionals
OR: odds ratio
PMWS: professional medical writing support
PRISMA: Preferred Reporting Items for Systematic Reviews and Meta-Analyses
RCT: randomized controlled trial
RR: relative risk
SD: standard deviation

## Declarations

### Ethics approval and consent to participate

Not applicable.

### Consent for publication

Not applicable

### Availability of data and material

Data sharing is not applicable to this article as no datasets were generated or analysed during the current study.

### Competing interests

Obaro Evuarherhe, Richard White and Christopher C Winchester are employees of Oxford PharmaGenesis, Oxford, UK. William Gattrell is an employee of Ipsen Pharma, Milton Park, UK. Christopher C Winchester and Richard White are directors of, and own shares in, Oxford PharmaGenesis Holdings Ltd.

### Funding

This study was funded by Oxford PharmaGenesis.

### Authors’ contributions

The study was conceived by CW and drafted by OE, WG, RW and CW.

## Acknowledgements

Results of this systematic review were presented as a poster at the 2018 European Meeting of ISMPP [36]. The authors would like to thank Charlotte Cookson and Gemma Carter for their assistance with reviewing this manuscript. The authors would also like to thank Dr Shruti Shah for provision of a full poster for one of the identified congress abstracts.

